# Plastid DNA sequences and oospore characters of some European species of *Tolypella* section *Tolypella* (Obtusifolia, Characeae) indicate a new cryptic *Tolypella* species from the Mediterranean island Sardinia

**DOI:** 10.1101/2022.11.11.516156

**Authors:** Anja Holzhausen, Petra Nowak, Andreas Ballot, Ralf Becker, Jasmina Gebert, Thomas Gregor, Kenneth G. Karol, Elisabeth Lambert, William Pérez, Uwe Raabe, Susanne C. Schneider, Nick Stewart, Klaus van de Weyer, Volker Wilde, Hendrik Schubert

## Abstract

In Europe, the genus *Tolypella* (Characeae) comprises four to eight *Tolypella* species in sections *Rothia* and *Tolypella* that have been distinguished by vegetative morphology and gametangial characters such as antheridial size and oospore cell wall ornamentation. However, morphological species differentiation is difficult in some cases due to overlapping and variable vegetative features, which in many cases are difficult to observe clearly.

To clarify the taxonomic status of the five European species of *Tolypella* in section *Tolypella*, sequence data of the plastid genes *atp*B, *rbc*L and *psb*C for *Tolypella glomerata* (Desv.) Leonh., *Tolypella hispanica* Allen, *Tolypella nidifica* (O.F. Müll.) A. Braun, *Tolypella normaniana* (Nordst.) Nordst. and *Tolypella salina* Cor. were combined with data on oospore morphology, including oospore wall ornamentation.

Gene sequence data identified five distinct clusters, but they differed from the morphologically identified five species. ‘*T. glomerata*’ consisted of some of the samples morphologically identified as *T. glomerata* and seven samples of *T. normaniana*, while the remaining *T. glomerata* samples clustered with specimens of unclear affiliation (“*Tolypella*. sp.”). ‘*T. hispanica* I’ consisted of samples from various locations, whereas “T. *hispanica* II” consisted of samples of *T. hispanica* from the Mediterranean island, Sardinia. The remaining cluster consisted of all the specimens that had been determined as *T. salina* or *T. nidifica* in addition to two specimens of *T. normaniana*. Oospore morphology was most clearly distinguishable for *T. glomerata*. Oospore characteristics for all other taxa were not as informative but showed some geographical and/or environmentally influenced differences, especially for *T. nidifica* and *T. salina*.

Our results suggest a significantly different taxonomy of *Tolypella* sect. *Tolypella* in which specimens normally identified as *T. glomerata* might be two different species, *T. glomerata* and an unidentified species; *T. nidifica* and *T. salina* are not separate species; *T. normaniana* is a diminutive variant of *T. nidifica* or *T. salina*; and *T. hispanica* comprises two different species, one from the Mediterranean island Sardnia.

## 1 Introduction

Charophytes, extant and fossil members of the order Charales plus the members of the extinct orders Sycidiales and Moellerinales (Schneider et al., 2015) are algae with a complex morphology, which are closely related to modern land plants (Nishiyama et al., 2018). Species delineation of charophytes is commonly based on morphological traits of the plant thallus, and accurate identification of charophyte species is important for understanding their diversity and for documenting changes in species distribution. Charophyte species identification is, however, hampered because of morphological plasticity influenced by abiotic factors. This specifically applies to the genus *Tolypella* A.Braun, where morphological characters are in some cases difficult to use because of (1) their small size and fragility, which often makes characters hard to observe; (2) phenotypic plasticity due to environmental influences such as water level and salinity (Lambert et al., 2013, Mouronval et al., 2015); and (3) their short vegetation periods that (a) often impede the use of characters derived from mature oospores (Wood, 1965) and (b) lead to fewer species collections due to their main development period being within a short time period that is easily missed. Some Characeae, particularly in the genus *Tolypella*, are ephemeral and seem to be rare. Most authors agree to split *Tolypella* into the two sections, *Tolypella* and *Rothia*, differentiated mainly by the shape of end cells (obtuse for *Tolypella* and acute for *Rothia*, (Krause, 1997) as well as habitat traits (Mouronval et al., 2015). Eight species of *Tolypella* have been described from Europe (Krause, 1997): five species are included in section *Tolypella* and include *Tolypella glomerata* (Desv.) Leonh., *Tolypella hispanica* Allen, *Tolypella nidifica* (O.F. Müll.) A. Braun, *Tolypella normaniana* (Nordst.) Nordst., and *Tolypella salina* Cor.. Species of the section *Rothia* are not considered in this study. There is no agreement about the taxonomic status of these species among different authors. For example, *T. nidifica* and *T. salina* were treated as distinct species by e.g. Krause (1997) or Mouronval et al. (2015) based on oospore features including ornamentation patterns since Corillion described *T. salina* as new species based on morphological and cytological criteria in 1960.

Of all the species in section *Tolypella*, only *T. hispanica* can be unambiguously differentiated, because it is dioecious, while all other *Tolypella* species are monoecious. Identification of the remaining four species has been mainly based on vegetative morphological traits, including oospore characteristics (Krause, 1997) and ecological ones (Wood, 1965). In order to aid species identification of the genus *Tolypella*, the additional use of oospore characteristics (e.g. length, number of striae and membrane ornamentation) has been suggested as occasionally useful (e.g. (Pérez et al., 2015). In addition, DNA barcoding, i. e. the use of short regions of DNA to identify species by assigning individuals to known taxa through comparison of their barcodes with a reference library, has become a popular means to improve species identification (McCourt et al., 1999, Sheth and Thaker, 2017). Moreover, DNA barcoding permits the identification of morphologically similar but genetically different (‘cryptic’) species, a common phenomenon for algae (Diaz-Tapia et al., 2018). Pérez et al. (2017) used the genes, *atp*B, *rbc*L and *psb*C successfully for species discrimination within the genus *Tolypella* in North America. Therefore, the same three plastid genes were also used in this study to investigate the species diversity of section Tolypella.

The aim of this study is to gain new insights into European *Tolypella* species by means of oospore characters combined with genetic data. For this, specimens of *T. glomerata, T. hispanica, T. normaniana, T. salina* and *T. nidifica* were examined, with the latter two included in such an attempt for the first time.

## 2 Materials and methods

Specimens identified as *T. glomerata, T. hispanica, T. nidifica, T. normaniana* and *T. salina* by means of vegetative characters (mostly antheridia sizes) were obtained from herbarium collections and from field collections by the authors for a total of 157 specimens. The collections span the period between 1871 and 2020 from locations in nine European countries (Austria, France, Germany, Great Britain, Greece, Ireland, Italy, the Netherlands, Norway, Portugal, Sweden) and from Chile in South America (**Table S1**). In addition, 40 specimens (Denmark, Italy, Germany, Great Britain, Greece, France, Norway, Portugal, Sweden) could not be unambiguously assigned morphologically to any recognized species and are referred to as *Tolypella* sp. throughout the manuscript.

Specimens of *T. nidifica* and *T. glomerata* from deep water sites were collected by diving. In shallow waters, samples were gathered by snorkeling or wading. Some specimens of *T. salina* (France) and *T. glomerata* (Germany) originated from germination experiments under laboratory conditions (e.g. Holzhausen 2016, Holzhausen et al. 2018). For all fresh material, oospores were harvested after release from cultured material in order to confidently assess oospore maturity. In addition, oospores of *T. nidifica* from Austria and Germany were collected from sediment samples. Fresh plant material was morphologically determined by the respective collector (**Table S1, Table S2**) based on species descriptions by various authors (e.g. (Corillion, 1975, Krause, 1997, Cirujano et al., 2008, Lambert et al., 2013). Herbarium specimens were sampled from collections deposited in the *Chara* herbarium (ROST-CHAR) of the University of Rostock (ROST), the New York Botanical Garden (NY), the Natural History Museum, University of Oslo (O) and private herbaria of the collectors (**Table S1**).

### 2.1 Genetic analyses

Dried plant material was obtained from a total of 193 individuals initially identified as *T. glomerata* (60 specimens), *T. hispanica* (15 specimens), *T. nidifica* (51 specimens), *T. salina* (50 specimens), *T. normaniana* (9 specimens), and 13 morphologically ambiguous *Tolypella* sp. Genetic data for the *atp*B, *psb*C and *rbc*L plastid genes presented in this study were obtained by three different working groups: A) the University of Rostock, B) the New York Botanical Garden and C) the Norwegian Institute for Water Research by the following methods.

Method A) Genomic DNA was extracted using the DNeasy Plant Mini Kit (Qiagen, Hilden, Germany), following the manufacturer’s protocol. Amplification of the plastid genes *rbc*L, *psb*C, and *atp*B was performed with 10 PCR cycles with one minute each of annealing at 94°C, extension at 55°C, and denaturation at 72°C, followed by one minute each for denaturation (94°C), annealing (52°C), and polymerisation (72°C) in 25 cycles. The amplified DNA was purified using the Biometra-innuPrep Gel ExtractionKit (Analytik Jena, Jena, Germany) according to the manufacturer’s instructions. Samples, were sequenced using a 3130×L GeneticAnalyzer (Applied Biosystems, NY, USA) with sequencing primers identical to the primers that were used for PCR reactions (**Table S3**). Obtained sequences were checked visually and aligned using BioEdit v.7.0.5.2 (Hall, 1999).

Method B) Genomic DNA was extracted using the Nucleon Phytopure DNA extraction kit (GE Healthcare Gio-Sciences, Pittsburgh, PA, USA, Pérez et al. 2014). The *atp*B, *psb*C and *rbc*L genes were amplified by a nested PCR reaction using either a PTC-200 DNAEngine® Thermal Cycler (Bio-Rad, Hercules, CA, USA) or a Mastercycler® pro S (Eppendorf AG, Hamburg, Germany). Initial PCR amplicons were generated through the following cycling program: initial denaturation at 95°C for 2 minutes; 35 cycles of 95°C for 15 seconds; 48°C for 15 seconds and 72°C for 30 seconds; and followed by a final extension at 72°C for five minutes. The resulting PCR product were used in a second round of PCR amplification to generate internal sequences using the same cycling program with the exception that the cycling was reduced to 30 cycles and the final extension time reduced to 30 seconds. Products from both PCR sets were sequenced at the University of Washington Genome Center (Seattle, WA, USA).

Method C) Genomic DNA from *Tolypella* material was isolated after (Schneider et al., 2016). PCR for the *rbc*l, *atp*B, and *psb*C genes was performed on a Bio-Rad CFX96 Real-Time PCR Detection System (Bio-Rad Laboratories, Oslo, Norway) using the iProof High-Fidelity PCR Kit (Bio-Rad Laboratories, Oslo, Norway). The following cycling protocol was used for all three genes: one cycle of 5 min at 94°C, and then 35 cycles each consisting of 10 s at 94°C, 20 s at 62°C, and 20 s at 72°C, followed by a final elongation step of 72°C for 5 min. PCR products were visualized by 1.5% agarose gel electrophoresis with GelRed staining (GelRed® Nucleic Acid Gel Stain, Biotium, Fremont, USA) and UV illumination. Amplification of the *rbc*L, *atp*B and *psb*C gene region was conducted using the primers listed in **Table S3**. In some cases, a nested PCR was conducted using the former PCR product as template and a second primer pair for a further PCR amplification. For sequencing the same primers and if necessary, intermediate primers were used (**Table S3**). Sequences were analyzed and aligned using Seqassem (version 04/2008) and Align (version 03/2007) MS Windows-based manual sequence alignment editor (SequentiX – DigitalDNA Processing, Klein Raden Germany) to obtain DNA sequence alignments, which were then corrected manually. For each PCR product, both strands were sequenced on an ABI 3730 Avant genetic analyzer using the BigDye terminator V.3.1 cycle sequencing kit (Biosystems, Applied Biosystems, Thermo Fisher Scientific Oslo, Norway) according to the manufacturer’s instructions.

Complete sequences of all three plastid genes could not be generated for every sample analyzed due to the differing qualities of the specimens (age, storing conditions, drying conditions after collection, etc.). Therefore, four different datasets were used for the phylogenetic analyses in order to obtain as much information as possible for all specimens. The first dataset included the three plastid gene sequences from 88 individuals, whereas three additional data sets were compiled for each plastid gene separately; the *atp*B dataset with a total of 1034 positions for 94 samples, the *psb*C dataset with a total of 1104 positions for 125 samples, and the *rbc*L dataset with a total of 1265 positions for 189 samples. In order to estimate evolutionary divergence, pair-wise uncorrected p-distances and the number of substitutions were conducted using MEGA version7 (Kumar et al., 2016). To uncover phylogenetic relationships, Bayesian inference (BI) and maximum likelihood (ML) trees were constructed, with evolutionary substitution models evaluated in MEGA v.7. The method selected the same best-fitting evolutionary model (GTR+I+Γ) for each of the four datasets. The ML algorithm was conducted in MEGA v.7 with 1000 bootstrap replicates. BI trees were performed with MrBayes 3.2.6 (e.g. Ronquist and Huelsenbeck, 2003) with a random starting tree and two independent runs of one cold and three heated chains, each using default parameters. Each analysis was run for 2 million generations with trees sampled every 1000 generations and the first 25% generations discarded as burn-in.

Due to small genetic distances among some taxa, intraspecific data often produce a variety of possible trees when using conventional tree building methods. In such cases, the relationship among taxa is best expressed by a network that is able to show alternative potential phylogenetic relationships within a single figure (Bandelt et al., 1999). Furthermore, networks allow the identification and illustration of ancestral alleles whereas phylogenetic trees treat all sequences as terminal taxa (Posada and Crandall, 2002). For that reason, Median-Joining (MJ) network analyses were performed using the PopART software v1.7.

### 2.2 Oospore analyses

The terminology of oospore characters in this study is based on Soulié-Märsche and García (2015). Descriptions of membrane ornamentation follow those of Frame (1977) and Urbaniak et al. (2012). Altogether, 712 mature oospores were harvested from herbarium specimens, fresh plant material and sediment samples. Oospores were collected from 12 specimens of *T. glomerata*, 19 of *T. nidifica*, 18 of *T. salina* and 10 of *T*. sp. The individual numbers of oospores examined and the appropriate pre- treatments are given in **Table S2**.

For stereomicroscopic analysis, oospores were photographed in lateral, apical, and basal views with a mounted camera. All oospores were stored in single wells of microtiter plates for potential re- examination or REM. Qualitative oospore characteristics that were examined included colour, shape and membrane ornamentation. To differentiate among the various brown hues of oospores, colour terms used in this study are clay brown, fawn brown, nut brown, chestnut brown, dark brown wine red and black brown (RAL Colour RAL Colour System). Quantitative characteristics included number of striae, expression of striae (prominence of striae), angle of striae to the longitudinal axis, oospore length and width, fossa width (average of 4 fossae), and the lengths of the outer lines of the pentagonal basal impression. Length measurements were calculated using ImageJ 1.50i. The ISI (isopolarity index; 100*(length/width)) was also calculated.

Scanning electron microscope (SEM) analysis of oospores were done at the Senckenberg Forschungsinstitut und Naturmuseum Frankfurt. Prior to SEM observations, few oospores were pre- cleaned (HoAc 5%) and all were dried by lyophilisation. Dry specimens were later sputter-coated with gold. SEM images of the surface of oospores and fossa walls (magnification 200-2500X) were produced with a JEOL JSM-6490 LV in high-vacuum mode by using secondary electrons and routinely applying an acceleration voltage of 20kV.

Oospore characters were tested for normality using the Shapiro-Wilk Test. Pairwise species tests were performed for different levels of analyses (species, country, region, type of location and plants) by the Kruskal-Wallis-Test (SPSS). P ≤ 0.05 was used as statistical significance for oospore analyses.

To identify (a) the correlation between oospore characteristics and regionality and (b) parameter combinations that might provide reliable discrimination, combined analyses of all quantitative and qualitative oospore features as well as their ratios, with the exception of the membrane ornamentation, were performed by nonmetric multivariate techniques using the Primer7 software package (Clarke and Gorley, 2015). Principal component analysis (PCA) was based on square root transformed data and Euclidean distance matrices.

## 3 Results

### 3.1 Vegetative morphology and gametangial characters

Fresh plant material was morphologically determined by the respective collector (**Table S1, Table S2**) based on species descriptions by various authors (Corillion, 1975, Wood, 1965, Krause, 1997, Cirujano et al., 2008, Lambert et al., 2013, Pérez et al., 2017, van de Weyer and Schmidt, 2018). According to those, all used *T. hispanica* were clearly identified by diocese, whereas *T. glomerata, T. salina* and *T. nidifica* were first determined by antheridial sizes, habitat occurrences and oospore ornamentation. Species that featured vegetative and antheridial characters of two species, e.g. *T. nidifica* and *T. salina* were determined as *T. sp*.

### 3.2 Genetic analyses

The phylogenetic analyses of the plastid gene sequences in each dataset recovered the 196 *Tolypella* individuals into five general clades that were denoted as, ‘*T. glomerata*’, ‘*T. nidifica*/*salina’*, ‘*Tolypella* sp.’, and two distinct ‘*T. hispanica*’ clades (‘*T. hispanica* I’ and ‘*T. hispanica* II’; **Figure 1, Table S4**). Supporting values for each cluster were given in the sections below. The gene sequence similarities of *Tolypella* individuals within each group were generally over 99.8 % (**Table 1**). However, support for their phylogenetic placements were unresolved or weakly to moderately supported in the single gene analyses. Phylogenetic resolution and support were greatest in the three-gene analyses. The results of the network analyses were comparable to the phylogenetic trees, with the same clusters recovered in both approaches (**Figure 1-2, Table S1, Figure S1-S3**). Results for the ML tree and the MJ network of concatenated gene sequences are shown in **Figure 1-2**. Complete trees and networks for single gene analyses are shown in the supplement (**Figures S1-S3**). The labels used to identify genetic groups correspond to those in **Table S1**.

**Figure 1.**
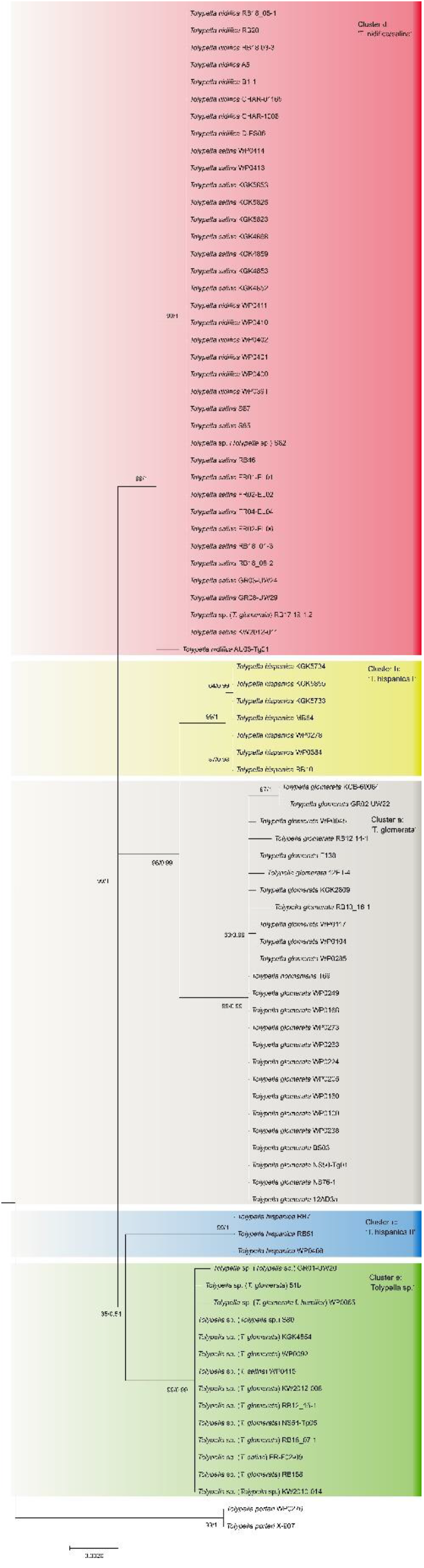
Maximum likelihood tree of genus *Tolypella* based on concatenated *atp*B, *psb*C, and *rbc*L sequence data. Phylogeny of Characeae based on combined *atp*B, *psb*C, and *rbc*L sequence data. Maximum likelihood tree with bootstrap values and posterior probabilities above branches (≥ 50 %).

**Table 1.** Estimates of evolutionary divergence based on concatenated dataset over sequence pairs between (black) and within (blue) main groups. Shown are the number of base differences (lower left) and the pairwise uncorrected p-distances (upper right) per sequence from averaging over all sequence pairs between and within groups.

|  | *T. glomerata* | *Tolypella* sp. | *T. nidifica/salina* | *T. hispanica* I | *T. hispanica* II |
| --- | --- | --- | --- | --- | --- |
| *T. glomerata* | 0.12% \| 3.85 | 27.73 | 26.77 | 18.00 | 31.92 |
| *Tolypella* sp. | 0.85% | 0.06% \| 2.00 | 19.40 | 24.00 | 23.40 |
| *T. nidifica/salina* | 0.82% | 0.59% | 0.01% \| 0.38 | 23.50 | 23.97 |
| *T. hispanica* I | 0.55% | 0.73% | 0.72% | 0.06% \| 2.00 | 30.00 |
| *T. hispanica* II | 1.01% | 0.73% | 0.72% | 0.92% | 0.00% \| 0.00 |

**Figure 2.**
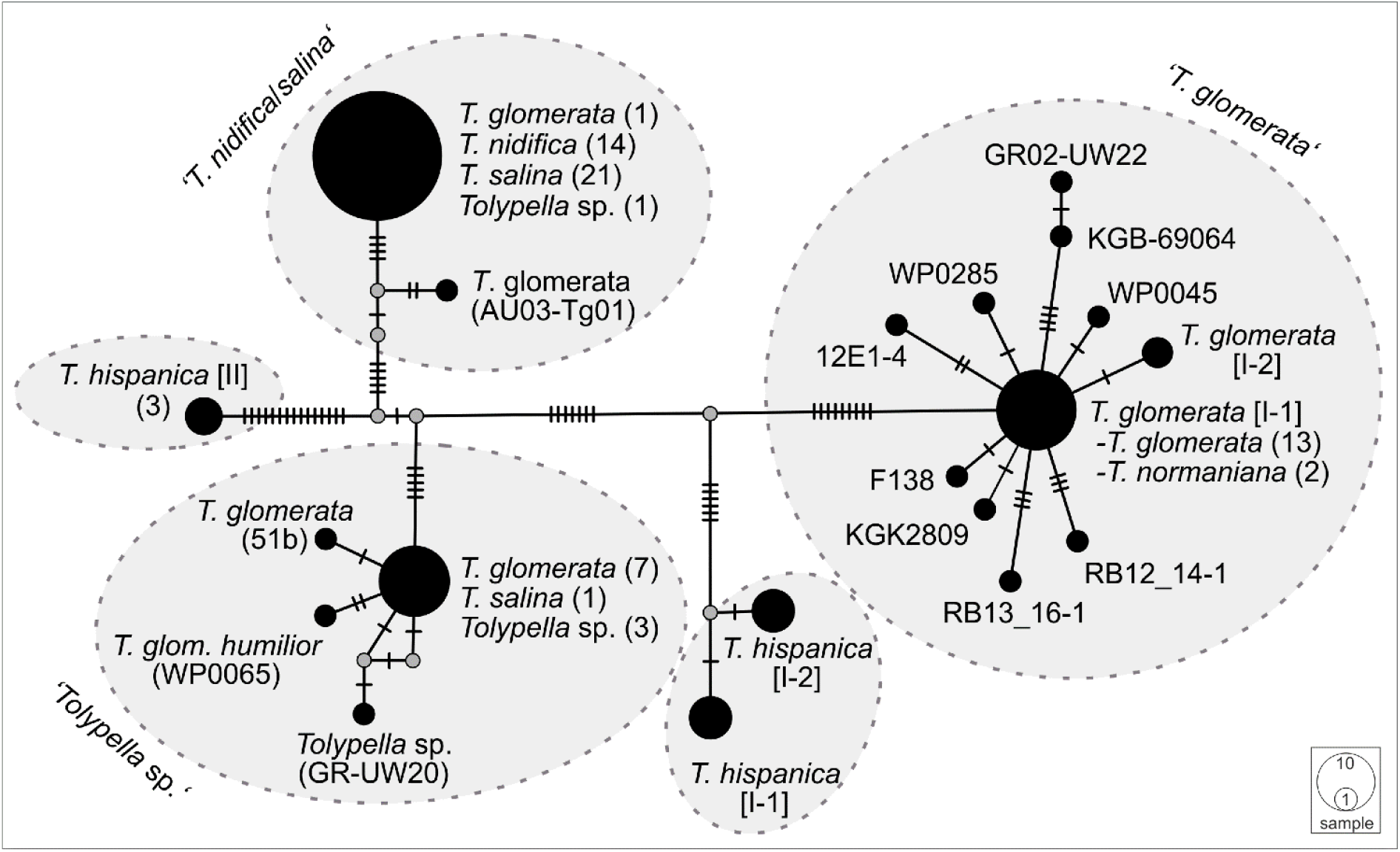
A Median Joining network of concatenated *atp*B, *psb*C, and *rbc*L sequences of sec. *Tolypella*. Circles represent haplotypes, with the size being proportional to their relative frequencies. The smallest circle corresponds to a single haplotype copy. A small black line at branches indicates one inferred mutational step. The small grey dot is a median vector and represents a possible extant unsampled haplotype or an extinct ancestral haplotype.

#### ‘T. glomerata’

A first group comprising mostly *T. glomerata* contained 49 individuals that represented *T. glomerata* (39 specimens), *T. normaniana* (7 specimens), and *T. sp*. (1 specimen). Specimens originated from nine European countries (France, Germany, Great Britain, Greece, Ireland, Italy, Netherlands, Norway, Sweden), and from the United States, Canada, and Chile. Analysing the concatenated sequences, ‘*T. glomerata*’ revealed ≥ 18 nucleotide differences to other clusters of sec. *Tolypella* (**Table 1**). With an average of about 3.85 bp substitutions, the genetic variability within the group was relatively high compared to the intragroup variability shown by the other species groups. However, differences in their sequence data were not regionally correlated; European and North American specimens showed identical sequences. In contrast, 8 substitutions were observed between samples collected in Greece (GR02-UW22) and Italy (RB13_16-1 and RB12_14-1).

#### ‘T. nidifica/salina’

A second cluster (labelled ‘*T. nidifica*/*salina*’) consisted of 110 individuals which have traditionally been assigned mainly to the species *T. nidifica* (44 specimens) and *T. salina* (50 specimens). Additionally, two *T. normaniana*, and 14 morphologically ambiguous *T*. sp. were found in this cluster (**Table S4**). They were collected in Austria, France, Germany, Greece, Italy, Norway and Sweden. Interestingly, the specimen originally determined as *T. glomerata* f. *littorea* from France (KGK4867) and two of nine sequenced *T. normaniana* (T70/T71) clustered within *T. nidifica/salina*. Sequence data for each of the *atpB, psbC* and *rbcL* genes could not be obtained for every specimen in this cluster. r*bc*L sequence data was obtained from 188 specimens, whereas sequences for *atpB* and *psbC* were obtained from 38 specimens (**Table S4**). Overall, however, there was little genetic variation within the ‘*T. nidifica*/*salina*’ cluster when comparing each of the gene sequences. Depending on the dataset, between 95.2 and 97.6 % of the analysed specimens had identical sequences. Minor genetic differences were observed in this group; two *T. sp*. collected from Italy (*rbc*L, **Figure S1**), and a *T. normaniana* from Norway (*psb*C, **Figure S3**) differed by a single nucleotide substitution each. Regional differences were not reflected in the sequence data with identical haplotypes throughout Europe. Consistent nucleotide differences were found only among two *T. sp*. collected in Austria (AU03-Tg01, AU03-Tg03).

#### ‘*Tolypella* sp’

A third cluster (labelled ‘*Tolypella* sp.’) consisted of 21 individuals which have been assigned as morphologically ambiguous specimens due to the presence of vegetative characters of more than one *Tolypella* species mentioned above. The specimens were originally determined as *T. glomerata* f. *humilor*. They were collected in Denmark, France, Germany, Great Britain, Greece, Italy, Norway, Portugal, and Sweden. ‘*Tolypella* sp.’ revealed unique sequence data and differed from ‘*T. nidifica*/*salina*’ and ‘*T. glomerata*’ by averaging 19.4 and 27.9 bp substitutions respectively (**Table 1**). Nucleotide differences within ‘*Tolypella* sp.’ ranged from 0 to 4 bp substitutions for concatenated sequences (mean 0.06 %, **Table 1, Figure 2**).

#### ‘T. hispanica’

Two clusters labelled as ‘*T. hispanica* I’ and ‘*T. hispanica* II’ consisted of ten and five individuals, respectively, which have traditionally been assigned to the dioecious *T. hispanica*. ‘*T. hispanica* I’ included ten individuals collected in France, Greece, Italy, and Algeria. The samples collected in France had two unique nucleotide substitutions for the combined sequences (0.06 %, **Table 1**). ‘*T. hispanica* I’ formed a strongly supported clade together with ‘*T. glomerata*’ (**Figure 1**). ‘*T. hispanica* II’ contained five individuals from four field collections in Italy, Sardinia that shared identical *rbc*L sequences whereas three individuals had identical sequences for all three genes (**Table 1**). In the ML analysis, however, ‘*T. hispanica* I’ was sister to ‘*Tolypella* sp.’ in a weakly supported relationship (**Figure 1**).

### 3.3 Oospore analyses

Differences in quantitative and qualitative oospore characters were considered with respect to species determined by either vegetative morphology or genetically determined species cluster. The results of oospores in species determined by vegetative morphology are summarized in **Table 2**.

**Table 2.**
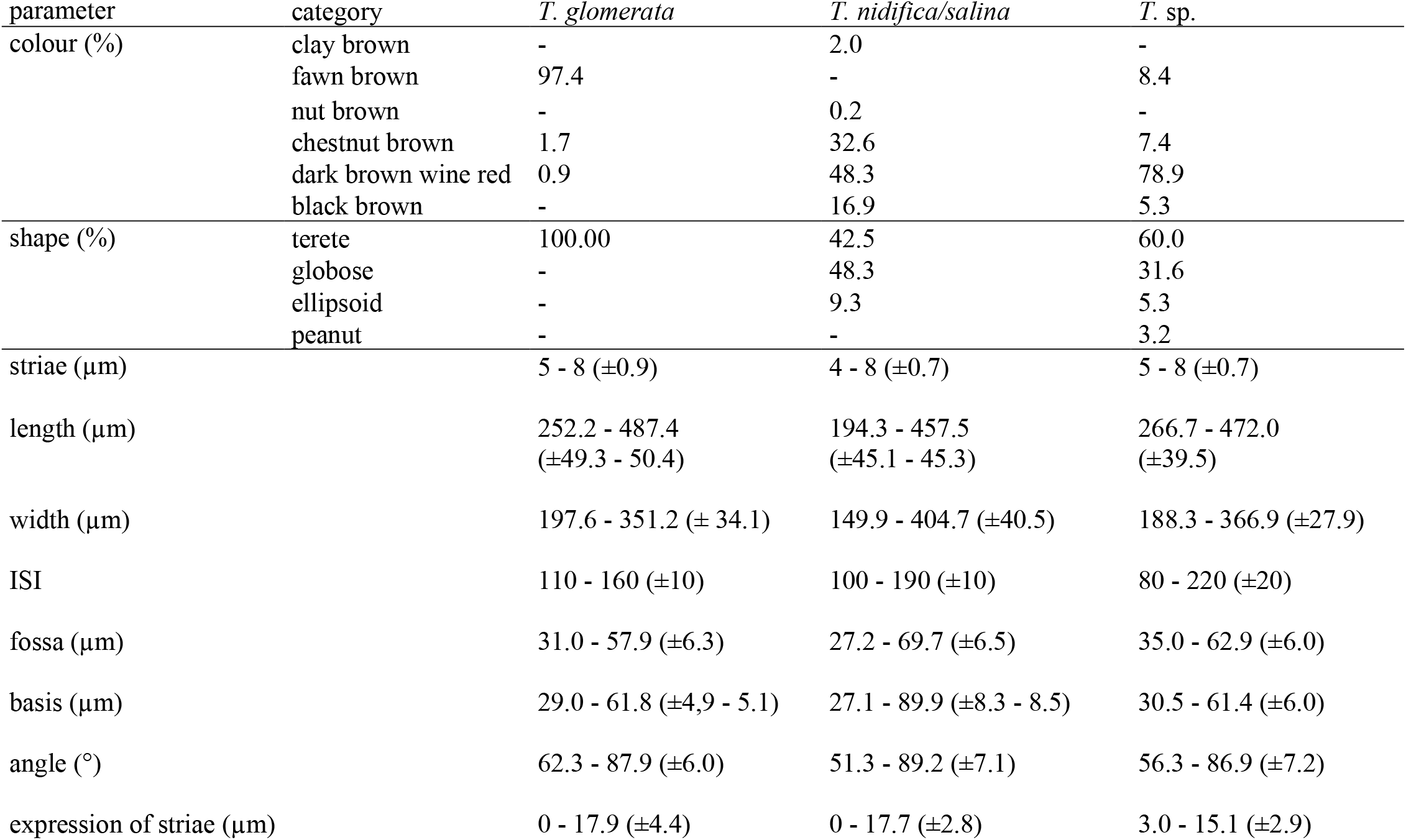
Oospore characteristics of *T. glomerata, T. nidifica/salina* and the morphologically unconclusive *T*. sp. Ranges of quantitative oospore features are given as min – max (± standard deviation).

| parameter | category | *T. glomerata* | *T. nidifica/salina* | *T. sp.* |
| --- | --- | --- | --- | --- |
| colour (%) | clay brown | - | 2.0 | - |
|  | fawn brown | 97.4 | - | 8.4 |
|  | nut brown | - | 0.2 | - |
|  | chestnut brown | 1.7 | 32.6 | 7.4 |
|  | dark brown wine red | 0.9 | 48.3 | 78.9 |
|  | black brown | - | 16.9 | 5.3 |
| shape (%) | terete | 100.00 | 42.5 | 60.0 |
|  | globose | - | 48.3 | 31.6 |
|  | ellipsoid | - | 9.3 | 5.3 |
|  | peanut | - | - | 3.2 |
| striae ($\mu\text{m}$) | | 5 - 8 ($\pm 0.9$) | 4 - 8 ($\pm 0.7$) | 5 - 8 ($\pm 0.7$) |
| length ($\mu\text{m}$) | | 252.2 - 487.4<br>($\pm 49.3$ - 50.4) | 194.3 - 457.5<br>($\pm 45.1$ - 45.3) | 266.7 - 472.0<br>($\pm 39.5$) |
| width ($\mu\text{m}$) | | 197.6 - 351.2 ($\pm 34.1$) | 149.9 - 404.7 ($\pm 40.5$) | 188.3 - 366.9 ($\pm 27.9$) |
| ISI | | 110 - 160 ($\pm 10$) | 100 - 190 ($\pm 10$) | 80 - 220 ($\pm 20$) |
| fossa ($\mu\text{m}$) | | 31.0 - 57.9 ($\pm 6.3$) | 27.2 - 69.7 ($\pm 6.5$) | 35.0 - 62.9 ($\pm 6.0$) |
| basis ($\mu\text{m}$) | | 29.0 - 61.8 ($\pm 4.9$ - 5.1) | 27.1 - 89.9 ($\pm 8.3$ - 8.5) | 30.5 - 61.4 ($\pm 6.0$) |
| angle ($^{\circ}$) | | 62.3 - 87.9 ($\pm 6.0$) | 51.3 - 89.2 ($\pm 7.1$) | 56.3 - 86.9 ($\pm 7.2$) |
| expression of striae ($\mu\text{m}$) | | 0 - 17.9 ($\pm 4.4$) | 0 - 17.7 ($\pm 2.8$) | 3.0 - 15.1 ($\pm 2.9$) |

#### T. glomerata

Oospores analysed in this study were usually fawn brown (97.4%), occasionally chestnut brown (1.7%) or dark brown wine red (0.9%), with an elongated rounded shape with 7-8 striae. The expression of striae is flat to prominent (0.0–17.9µm). Oospores showed lengths of 252.2 to 487.4 µm (±49.8-50µm), widths from 197.6 to 351.2 µm (±33.5-34.1µm), a mean fossa width of 31.0 to 57.9 µm, mean lengths of the outer basal impression from 29.0 to 61.8 µm and an ISI of 110–160. All oospores exhibited a reticulate ornamentation in varying expression and size (**Figure 3, Table S2**).

**Figure 3.**
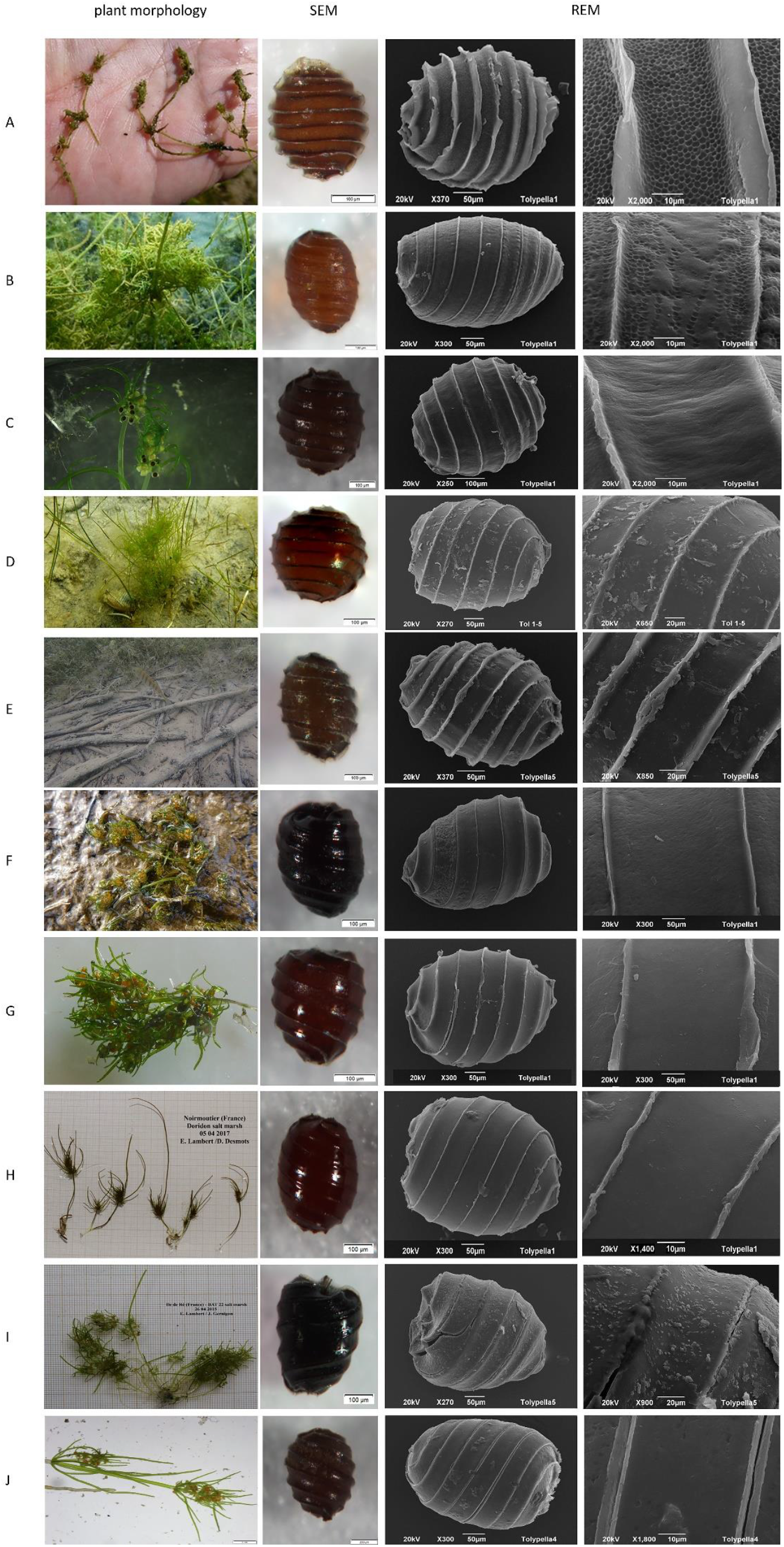
Habitus, SEM and REM of ‘*T. glomerata’*, ‘*T. nidifica’*, ‘*T. salina’* and ‘*T*.*sp*’. A- ‘*T*.*glomerata*’ RB13_16 (Italy, Cabras) with fully reticulate ornamentation pattern, B- ‘*T*.*glomerata*’ BS-Tol (Germany, Borkener See) with partially reticulate ornamentation of oospores. C-’*T. nidifica*’ TN3-1 (Germany, Fehmarn), D-’*T. nidifica*’ Tol04 (Germany, Lehmkenhafen), E-’*T. nidifica*’ Tol7 (Austria, Apetlon Badesee), F-’*T. nidifica*’ RB18_12 (Italy, Pittulongu). G-’*T. salina*’ RB18-01 (Italy, Pittulongu), H-’*T. salina*’ FR-EL/Sal1-07 (France, Île de Noirmoutiers), I-’*T*.*salina*’ FR-TS 687-01 (France, Île de Ré). J-’*T. sp*.’ FR-F02 (France, Kermadec).

#### T. nidifica/salina

Oospores of *T. nidifica/salina* were mainly dark brown wine red (48.3%), chestnut brown (32.6%) or black brown (16.7%) with a terete or broad rounded/globose shape and a flattened base. Oospores showed (4-)5-7(- 8) striae that were flat and prominent (0.0–17.7 µm), oospore lengths of 194.3– 457.5 µm (±45.2µm), oospore widths of 149.9–404.7 µm (±40.4µm), mean fossa width of 27.2 – 69.7 µm and an outer mean basal impression length of 27.1 to 89.9 µm. The calculated ISIs ranges between 100 and 190. Ornamentation patterns of *T. nidifica/salina* were highly variable, from smooth to smooth with some pustules or with fine linear structures (**Figure 3, Table S2**).

#### *Tolypella* sp

The oospores of the morphologically ambiguous specimens were dark brown wine red (78.9 %), occasionally fawn brown (8.4%), chestnut brown (7.4%) or black brown (5.3%) with a broad range of shape variations (ellipsoid, elongate rounded, broad rounded/globose, or peanut- shaped). Oospores showed (5) 6–7 (-8) striae with widths between 3.0–11.2 µm, oospore lengths of 266.7– 472.0 µm (±38.7µm) and widths between 195.6–346.7 µm (±27.9 µm). Fossae ranged from 37.2 µm to 62.2 µm and outer basal impression lengths from 33.1 up to 57.9 µm. Calculated ISIs ranges from 80 to 190 (-220). The membrane of *T. sp*. Oospores was smooth to smooth with some pustules or with fine linear structures ((**Figure 3, Table S2**).

The analysis of species-related oospore characteristics shows that variations, especially with regard to the features colour, shape and length exist within each species ((**Figure 3, Table 2, Table S2**). Especially for *T. glomerata*, large discrepancies between oospores along a geographical gradient / between populations could be observed (**Figure 4**). The PCA shows that the two axis explain 64,8% of the cumulative variation of oospores (eigenvalue 1 = 6,72, eigenvalue 2 = 3; **Table S5**). The first component is determined by the characters length and width and the ratios of length/angle and width/angle, whereas the second component is defined by the striae and width/fossa ratio. Depending on the level of analysis, significant intraspecific differences between countries, regions, type of locations and plants can be detected. Oospores from Germany (length: 335µm – 487µm, width: 255µm – 351µm) and Austria (length: 378µm – 438µm, width: 241µm – 313µm) were significantly larger and wider than those from Greece (length: 294µm – 326µm, width: 197µm – 221µm; p ≤ .001) and Italy (length: 252µm – 362µm, width: 203-µm – 274µm; p ≤ .001). No significant differences could be detected between oospores from Germany and Austria or between those from Italy and Greece.

**Figure 4.**
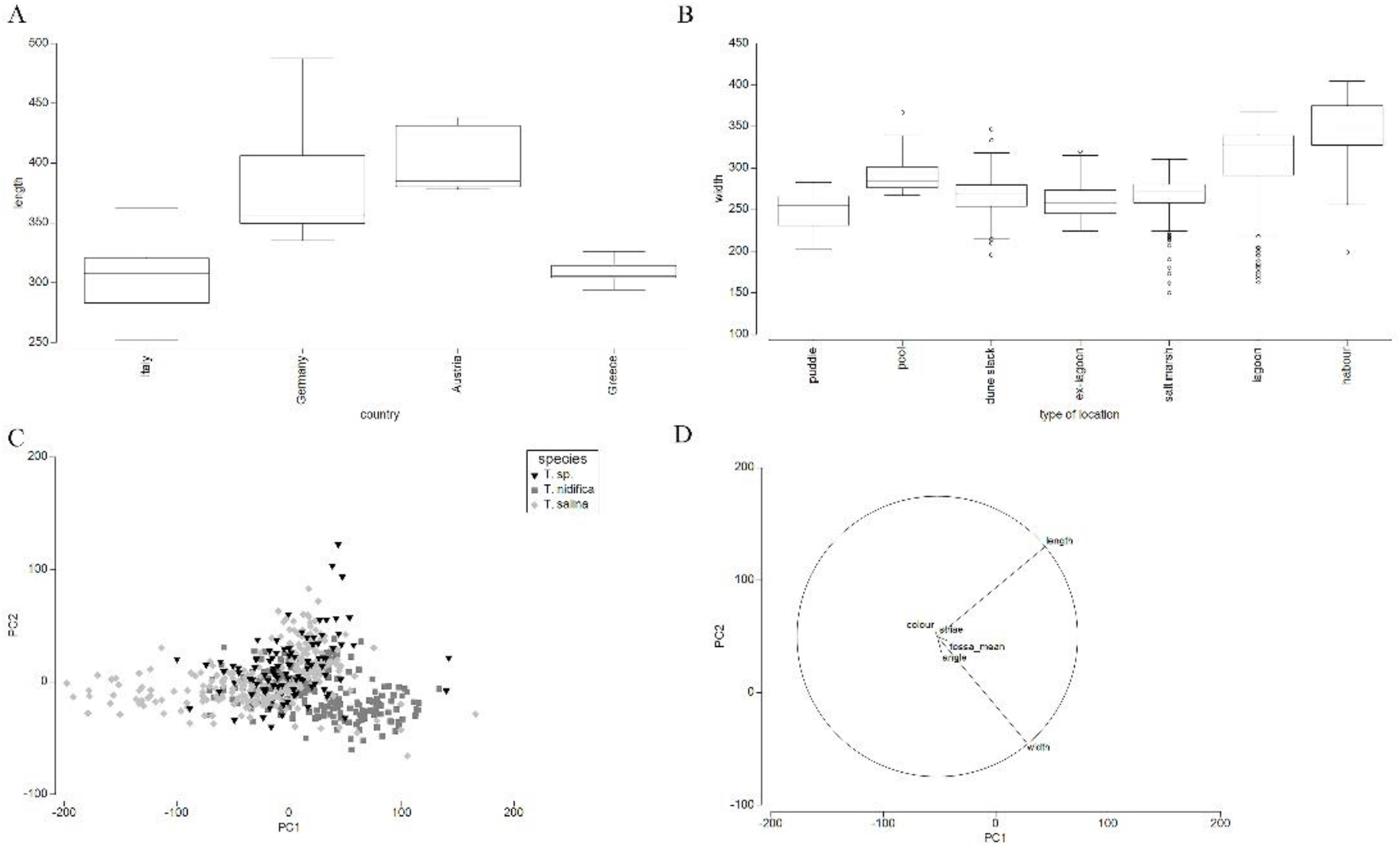
Analyses of oospore features. A- Box-Plot of *T. glomerata* depending on the country. B- Box-Plot of locally separated *T. glomerata* populations. C - PCA of vegetative determined *T. nidifica, T. salina* and *T*. sp. Oospores depending on the country. Included were 565 oospores from Germany, France and Italy. D – Characters of vegetative determined *T. nidifica, T. salina* and *T*. sp.

Given the vegetative determination in *T. nidifica, T. salina* and *T*. sp, significant differences could be obtained for the oospore characters shape (p ≤ .006), length (p ≤ .005), fossa (p ≤ .005) and width (p≤ .005) between *T. nidifica* and *T. salina. Tolypella sp*. could be separated from *T. nidifica* by the width, fossa (p ≤ .006) or shape (p ≤ .017) whereas *T. sp*. could be differentiated by oospore length (p≤ .005) and shape (p ≤ .006) from *T. salina* although overlapping areas exist.

Interestingly, depending on the type of location, oospores of permanent water bodies (lagoons, harbours and lakes) and pools are wider than oospores from temporary water bodies (**Figure 4**).

## 4 Discussion

The results of genetic analyses for European *T. glomerata* and *T. salina/nidifica* specimens can be confirmed only partly by oospore analyses. However, few examples showed that identified genetic differences could be confirmed by oospore features, especially wall ornamentation pattern. The results are only partially consistent with the current phenetic taxonomic concept (Groves and Bullock-Webster, 1920-1924, Corillion, 1975, Krause, 1997, Cirujano et al., 2008, Mouronval et al., 2015). Both analyses can confirm differences of unambiguous *Tolypella* specimens to *T. glomerata* and *T. nidifica/salina*. Based on genetics they are located within the *‘Tolypella* sp.’ Cluster, whereas oospore analyses revealed significant differences in e.g. length and width. In contrast to the sequence data, significant differences could be detected between oospore lengths and widths of *T. nidifica* and *T. salina*. But these are mainly caused by local separations. However, Italian *T. nidifica* did not differ from French *T. salina* or Italian/French *T*.sp. This is in strong accordance with the results of sequence data. Differences in oospore ornamentation patterns were not reflected by sequence data of plastid genes.

*Tolypella hispanica* is the only dioecious species in the section *Tolypella* and, by including sexuality as a taxonomically relevant parameter at species level, easily distinguished from all other European species. Phylogenetic data of this study revealed, that this taxon consists of two well-separated lineages, representing two cryptic species. One lineage (‘*T. hispanica* I’), in agreement with the results of Peréz et al. (2016), was related to *T. glomerata*. The second lineage (‘*T. hispanica* II’), identified for the first time in this study, was located very distant from ‘*T. hispanica* I’ and nearer to the cluster of ‘*Tolypella* sp.’ and ‘*T. nidifica/salina*’. Several studies demonstrated that sex separation occurred independently in various groups of the Characeae (Proctor, 1980, Meiers et al., 1999, Pérez et al., 2017). The results of our analyses revealed the existence of two cryptic species, being united within the recent taxon *T. hispanica*. The existence of cryptic species is very common for about half of all marine eukaryotic organisms such as Rhodophyta (e.g. Payo et al., 2013), Chlorophyta (Cimino and Delwiche, 2002, Irisarri et al., 2021) or Phaeophyceae (Poong et al., 2013). In order to get a robust description of morphological characters for discrimination between the two dioecious taxa, detailed analysis of a larger number of samples needs to be done. Although nature conservation is extremely important especially for Mediterranean islands such as Sardinia, Charophytes are not included in Sardinian conservation programmes so far. Becker (2019) highlighted the presence of 26 different charophyte species in Sardinia with respect to their habitat preference. Moreover, he suggested four different action plans for (I) Characeae of lagoons, temporary brackish pools, salt marshes and estuaries, (II) *Nitella* species of temporary freshwater ponds and estuaries, (III) *Chara connivens* in temporary ponds and lakes and (IV) *Chara* species of running waters in calcareous regions and water reservoirs to counteract the loss of species and habitats, including the new cryptic species of *T. hispanica*.

For *T. glomerata*, a broad morphological variability within European specimens, comparable to the results published by (Pérez et al., 2014) for North American specimens, was observed in this study. The shape of the whorls ranged from very compressed with short or long sterile branchlets to whorls which rather appear looser with long sterile branchlets (spike-like). This broad morphological variability is also reflected by oospore characters, exhibiting large regional differences (**Figure 3**). On the other hand, only small differences among gene sequences of European *T. glomerata* and those from North and South America could be detected (**Figure 2**). The results of this study show that even with a broader sampling range the ‘*T. glomerata*’ cluster remains stable. For North American specimen, (Pérez et al., 2015) found a reticulate oospore ornamentation for *T. glomerata*, the most useful character for the distinction between *T. porteri* and *T. glomerata*. Antheridia size, on the other hand, seems not to be a suitable character for discrimination between *T. glomerata* and *T. nidifica*/*salina*. For example, eight individuals have to be identified as *T. glomerata* using antherida sizes but were genetically determined as *T. nidifica*/*salina*. Although for *T. glomerata* smaller antheridia sizes in diameter (220–450µm; (Corillion, 1960, Krause, 1997) are reported than for *T. salina* (450–625µm (–1000µm) (Corillion, 1960, Krause, 1997, Cirujano et al., 2008, Lambert et al., 2013) and *T. nidifica* (450–550µm; (Krause, 1997, Urbaniak, 2003). This character is often used for discrimination between *T. glomerata* and *T. nidifica* in the field, but seems to be influenced by environmental conditions as reflected by a North-South gradient, resulting in a broad and overlapping size range for these taxa. Clear and unambiguous distinction between *T. salina* and *T. nidifica* could not be achieved by this study. Neither by means of genetic data nor by oospore morphology and ornamentation. The comparison of French and Italian *T. salina* (Lambert et al., 2013, Becker, 2019) with those of the Iberian Peninsula (Cirujano et al., 2008, Cirujano Bracamonte et al., 2013) depicts a large morphological variability, probably caused by environmental conditions. A correlation between habitat salinity and phenotypic plasticity/fructification has already been published for several halophytic charophytes (e.g. Winter and Kirst, 1991, Bonis et al., 1993). Both *T. salina* phenotypes, (1) smaller plants with fewer and shorter fertile branchlets (Cirujano et al., 2008) and (2) bigger ones with a higher number and longer fertile branchlets and internodes (Lambert et al., 2013), have been identified in this study for France as well as Italy (**Figure 3**). Those interannual morphological variability is caused by environmental variability and well known for Characeae.

A similar large morphological variability appeared within *T. nidifica*. Specimens with very compact and compressed whorls, long branchlets and long internodes, as well as specimens with less compact and compressed whorls and shorter branchlets were observed (**Figure 3**). The same applies for oospore morphometry, also exhibiting large variability without being reflected by genetic differences of the standard marker genes investigated here. Morphometric oospore characters exhibited site-specific and location-specific differences, but did not allow for discrimination between the two taxa. As for vegetative characters, the reason for this observed large variability might be habitat conditions such as the (soil) salinity. Oospores from puddles could be clearly differentiated from higher saline locations such as salt marshes, lagoons or harbours. The Italian sites exhibited a salinity between 1.2 and 21.6g/L (Becker, 2019) the salinities of the French salt marshes ranged from 2.2 to up to 250g/L (Lambert et al., 2013) with large seasonal changes whereas for the German Lehmkenhafen (2020) salinity is less fluctuating around 12. Seasonal changes are mainly caused by drought and re-wetting of temporal ponds or puddles.

*T. salina* was described by Corillion (1960) who gives the differences from *T. nidifica* as the lower number of striae of the oospore (mostly 6 vs. 8) and smaller oospores (length 273–366 µm vs. 400–475 µm; width 258–312 µm vs. 350–450 µm). These differences could not be corroborated by us (**Table 2**).

Until now, besides the number of chromosomes (50 for *T. salina* and 20-42 for *T. nidifica*, (Corillion, 1960, Guerlesquin, 1967), the membrane ornamentation was found in this study as most reliable character although no clear distinction is possible. Whereas the oospores of *T. salina* showed a smooth ornamentation, excepting a few specimens with only few pustules, which is only partially in accordance with different authors (Corillion, 1975, Urbaniak et al., 2012), the membrane of *T. nidifica* exhibited in most cases pustules or linear structures, only few specimens revealed smooth oospores. However, the number of oospores available for examination was rather low and, moreover, this result partly contradicts existing literature. Nordstedt (1889) described the oospore membrane of *T. nidifica* as smooth, Wood (1965), Ray et al. (2001) and Urbaniak et al. (2012) found a pit-like ornamentation for *T. nidifica* which ‘varied among populations’ and Corillion (1975) described both expressions. The results of this study also showed transitions between both ornamentation types which should be investigated in more detail as well as in corelation to the maturity status of oospores which was shown by Casanova (1991) for *Nitella* oospores.

In addition to overlapping morphological plant features, determinations may be hampered by the existence of intermediary forms between *T. glomerata* and *T. nidifica* as described for the French population from Herault (Hy, 1913, Corillion, 1957). So as for the vegetative characters, habitat-specific effects on oospore ornamentation needs to be investigated in more detail by acclimation experiments before a definite conclusion about the reliability of ornamentation pattern for species delineation can be made. However, a distinct genetic entity, until now represented just by one specimen, was detected. This specimen originated from a brackish lake near Apetlon in Austria/Burgenland (AU03-Tg01) and is the first record of *T. nidifica*/*salina* for Austria and should be investigated in more detail.

Consequently, a final conclusion about the taxonomic status of *T. nidifica*/*salina* cannot be made irrespective of the observed differences in ornamentation pattern. With respect to lacking genetic differences, Pérez et al. (2017) have shown that analyses based on ribosomal gene sequences support chloroplast data but are not reliable for discrimination between uncertain taxa.

Nevertheless, both analyses could be considered as appropriate, and imply possibilities for further investigations of the species status of these European taxa such as analyses of geographically isolated *Tolypella* populations on the basis of Simple Sequence repeats as microsatellite studies have shown for the genus *Chara* (Schaible et al., 2012, Noedoost et al., 2015). NGS- techniques, multi-omics approaches including proteomics could be carried out to examine smallest genetic differences as shown for *Nitellopsis obtusa* (Sleith and Karol, 2021).

## Conclusion

This study showed that the combination of oospore morphology and sequence data are only partially consistent. Sequence data confirmed the taxonomic status of *T. glomerata* and *T. hispanica*. Besides this, a second dioecious *T. hispanica* lineage can be identified.

Moreover, although *T. nidifica* and *T. salina* could not be separated by sequence data (‘*T. nidifica/salina’)* and transitions in oospore ornamentation exist, this study reveals significant differences in oospore length and widths that are mainly caused by local differences. These results indicate that environmental influences are in correlation with oospore morphology. The species rank of *T. normaniana* could not be confirmed by genetic results. Those individuals clustered within ‘T. *glomerata*’ and ‘T.*nidifica*/*salina*’. Furthermore, the sequence data revealed a new genetic entity, currently named as *T*.*sp*. A final decision about the taxonomic status of *T. nidifica*/*salina* and *T*.sp could not be done on the basis of these results. Nevertheless, all analyses could be considered useful, and imply possibilities for further investigations of the species status of these European taxa.

## 6 Author Contributions

Conceptualization - A.H.; PN methodology – A.H., A.B., S.C.S., T.G., P.N., K.G.K., W.P., V.W.; analysis/investigation: A.H., A.B., S.C.S., T.G., P.N., K.G.K., W.P., V.W., E.L., U.R., R.B., K. v. d. W.; resources – R.B., E.L., U.R., N.S., K. v. d. W., H.S.; original draft preparation - A.H.; review and editing – all authors, visualization - A.H., P.N.

## 7 Acknowledgments

The authors want to thank Peter Gisch from the Biological Station Neusiedler See for the provision of physico-chemical data. They are also grateful to Didier Desmots from the National Natural Reserve of the Müllembourg salt marshes, Isle of Noirmoutier, Barbara Tuner and Karl-Georg Bernhardt for collecting specimens and environmental data. Moreover, the authors are thankful to Claudia Lott (University of Rostock) for preparing genetic analyses in Rostock and Arne Schoor for laboratory and technical assistance. The laboratory equipment was partly supported by the European Fund for Regional Development (EFRD).

## 8 Supplementary Material

**Supplementary Figure 1**. Phylogeny of Characeae based on *rbc*L sequence data. (A) Median Joining network of *rbc*L sequences of *Tolypella*. (B) Maximum likelihood tree of genus *Tolypella* based on *rbc*L sequence data with bootstrap values and posterior probabilities above branches (≥ 50 %).

**Supplementary Figure 2**. Phylogeny of Characeae based on *atp*B sequence data. (A) Median Joining network of *atp*B sequences of *Tolypella*. (B) Maximum likelihood tree of genus *Tolypella* based on *atp*B sequence data with bootstrap values and posterior probabilities above branches (≥ 50 %).

**Supplementary Figure 3**. Phylogeny of Characeae based on *psb*C sequence data. (A) Median Joining network of *psb*C sequences of *Tolypella*. (B) Maximum likelihood tree of genus *Tolypella* based on *psb*C sequence data with bootstrap values and posterior probabilities above branches (≥ 50 %).

**Supplementary Table S1**. List of the 195 *Tolypella* samples used for oospores and genetic analyses in the present study. The columns “oospore ornamentation and “Genetics” columns indicate the results of oospore ornamental analysis as well as the DNA sequencing results. Detailed information is given in Table S2 (oospore analyses) and S4 (genetic analyses). The last columns show the accession numbers, with n.n. in case the genetic marker was not recovered. (Tglo=*T. glomerata*, Tni=*T. nidifica*, Tsal=*T. salina*, Tn/s=*T. nidifica*/*salina*, Thi=*T. hispanica*, Tsp=*T*. sp.). References : A - Nowak and Schubert 2019, B - Peréz et al. 2014, C - McCourt et al. 1999, D - Perez et al. 2016; storage: 1 - Herbarium University Rostock, 2 New York Botanical Garden Steere herbarium, 3 - Vascular Plant Herbarium, National History Museum Oslo, 4 - Herbarium Vienna, 5 - Friener Herbarium, Butler University, 6 - United States National Herbarium). (Excel Data Sheet)

**Supplementary Table S2**. List of oospores used for SEM and REM analyses including oospore morphology and ornamentation and the irresprective genetic cluster. Kind of material: F - fresh plant material, H - herbarium specimen, S - sediment sample, G - germination experiment. (Excel Data Sheet)

**Supplementary Table S3**. Oligonucleotide primers used in this study. “Method” refers to the method used for DNA-sequencing described in genetic analysis subsection in Materials and Methods. (Excel Data Sheet).

**Supplementary Table S4**. List of 195 *Tolypella* specimens used for genetic analyses in the present study. The first column indicates the genetic identification of the sample, while the second column contains the morphological determination according to the plant characteristics. “Method” refers to the method used for DNA-sequencing described in genetic analysis subsection in Materials and Methods. The last four colums display the genetic cluster observed by analysing single gene sequence data and a combined data set. Cluster names correspond to Figure S1 (*rbc*L), Figure S2 (*atp*B) and Figure S3 (*psb*C), and Figure 1 and Figure 2 (*atp*B + *rbc*L + *psb*C).

**Supplementary Table S5**. List of eigenvalues and eigenvectors resulted from the principal component analysis of all *Tolypella* samples including absolute oospore values and their ratios.

## 9 Data Availability Statement

Data are already published and publicly available, with those items properly cited in this submission:

1. McCourt RM, Karol KG, Casanova, MT, Feist, M. 1999. Monophyly of genera and species of Characeae based on *rbc*L sequences, with special reference to Australian and European Lychnothamnus barbatus (Characeae: Charophyceae). Aust. J. Bot. 47, 361- 369.
2. Pérez W, Hall JD, McCourt RM, Karol KG. 2014. Phylogeny of North American Tolypella (Charophyceae, Charophyta) based on plastid DNA sequences with a description of Tolypella ramosissima sp. nov. J Phycol 50: 776-789.
3. Pérez W, Casanova MT, Hall JD, McCourt RM, Karol KG. 2017. Phylogenetic congruence of ribosomal operon and plastid gene sequences for the Characeae with an emphasis on *Tolypella* (Characeae, Charophyceae). Phycologia 56(2): 230-237.
4. Nowak, P, Schubert H. 2019. Genetic variability of charophyte algae in Baltic Sea area. Botanica Marina 62(1): 75-82.

New Data Sets will be archived in permanent repositories (GenBank) and are provided with requestes Accession Numbers.

## References

Bandelt, H.-J., Forster, P. & Röhl, A. 1999. Median-Joining Networks for Inferring Intraspecific Phylogenies. Mol. Biol. Evol., 16, 37–48.

Becker, R. 2019. The Characeae (Charales, Charophyceae) of Sardinia (Italy): habitats, distribution and conservation. WEBBIA.

Bonis, A., Grillas, P., Van Wijck, C. & Lepart, J. 1993. The effect of salinity on the reproduction of coastal submerged macrophytes in experimental communities. Journal of Vegetation Science, 4, 461–468.

Casanova, M. T. 1991. An Sem study of developmental variation in oospore wall ornamentation of three Nitella species (Charophyta) in Australia. Phycologia, 30, 237–242.

Cimino, M. T. & Delwiche, C. F. 2002. MOLECULAR AND MORPHOLOGICAL DATA IDENTIFY A CRYPTIC SPECIES COMPLEX IN ENDOPHYTIC MEMBERS OF THE GENUS COLEOCHAETE BRÉB.(CHAROPHYTA: COLEOCHAETACEAE). Journal of Phycology, 38, 1213–1221.

Cirujano Bracamonte, S., Guerrero Maldonado, N. & García Murillo, P. 2013. The genus Tolypella A Braun A Braun in the Iberian Peninsula. Acta Botanica Gallica, 160, 121–129.

Cirujano, S., Chambra, J., Sánchez Castillo, P. M., Meco, A. & Flor Arnau, N. 2008. Flora ibérica Algas Continentales. Carófitos (Characeae), Madrid, Real Jardín Botánico, CSIC.

Clarke, K. R. & Gorley, R. N. 2015. Getting started with PRIMER v7. PRIMER-E Ltd.

Corillion, R. 1957. Les Charophycees de France et d’ Europe Occidentale. Bull. Soc. Sci., Bretagne, 32, 499.

Corillion, R. 1960. Tolypella salina sp. nov., Chorophycee nouvelle des morois de Croix-de-Vie (Vendee).

Corillion, R. 1975. Flore des charophytes du Massif armoricain et des contrées voisines d’Europe occidentale, Paris, France: Jouve.

Diaz-Tapia, P., Maggs, C. A., Macaya, E. C. & Verbruggen, H. 2018. Widely distributed red algae often represent hidden introductions, complexes of cryptic species or species with strong phylogeographic structure. J Phycol, 54, 829–839.

Frame, P. W. 1977. Fine Structural Studies of Oospore Ornamentation and Bulbil Development in Charophytes. phD, University of Toronto.

Groves, J. & Bullock-Webster, G. R. 1920–1924. The British Charophyta, London, The Ray society.

Guerlesquin, M. 1967. Recherches caryotypiques et cytotaxinomiques chez les Charophycées d’Europe occidentale et d’Afrique du Nord Thèse Université de Toulouse.

Hall, T. A. 1999. BioEdit: a user-friendly biological sequence alignment editor and analysis program for Windows 95/98/NT. Nucleic acids symposium series. Oxford University Press.

Holzhausen, A. (2016). Resilienz aquatischer Ökosysteme -Vitalitätsabschätzungen von Diasporenbanken, dissertation. (Rostock: University Rostock) doi: 10.18453/rosdok_id00002020.

Holzhausen, A., Porsche, C., and Schubert, H. (2017). Viability assessment and estimation of the germination potential of charophyte oospores: testing for site and species specificity. Bot. Lett. 165, 147–158. doi: 10.1080/23818107.2017.1393460

Hy, M. 1913. Les Characées de France. Bulletin de la Société Botanique de France, 60, 47.

Irisarri, I., Darienko, T., Proschold, T., Furst-Jansen, J. M. R., Jamy, M. & De Vries, J. 2021. Unexpected cryptic species among streptophyte algae most distant to land plants. Proc Biol Sci, 288, 20212168.

Krause, W. 1997. Charales (Charophyceae). In: Ettl, H., Gärtner, G., Heynig, H. & Mollenhauer, D. (eds.) Süßwasserflora von Mitteleuropa. Gustav Fischer Verlag.

Kumar, S., Stecher, G. & Tamura, K. 2016. MEGA7: Molecular Evolutionary Genetics Analysis Version 7.0 for Bigger Datasets. Mol Biol Evol, 33, 1870–4.

Lambert, E., Desmots, D., Le Bail, J., Mouronval, J.-B. & Felzines, J.-C. 2013. Tolypella salina R. Cor. on the French Atlantic coast: Biology and ecology. Acta Botanica Gallica, 160, 107–119.

Mccourt, R. M., Karol, K. G., Casanova, M. T. & Feist, M. 1999. Monophyly of Genera and Species of Characeae based on rbcL Sequences, with Special Reference to Australian and European Lychnothamnus barbatus (Characeae: Charophyceae). Aust. J. Bot., 47, 361–369.

Meiers, S. T., Proctor, V. W. & Chapman, R. L. 1999. Phylogeny and Biogeography of Chara (Charophyta) Inferred from 18S rDNA Sequences. Aust. J. Bot., 47, 347–360.

Mouronval, J.-B., Baudouin, S., Borel, N., Soulié-Märsche, I., Klesczewski, M. & Grillas, P. 2015. Guide des Characées de France Mediterranéenne, Office National de la Chasse et de la Faune Sauvage.

Nishiyama, T., Sakayama, H., De Vries, J., Buschmann, H., Saint-Marcoux, D., Ullrich, K. K., Haas, F. B., Vanderstraeten, L., Becker, D., Lang, D., Vosolsobe, S., Rombauts, S., Wilhelmsson, P. K. I., Janitza, P., Kern, R., Heyl, A., Rümpler, F., Calderón Villalobos, L. I. A., Clay, J. M., Skokan, R., Toyoda, A., Suzuki, Y., Kagoshima, H., Schijlen, E., Tajeshwar, N., Catarino, B., Hetherington, A. J., Saltykova, A., Bonnot, C., Breuninger, H., Symeonidi, A., Radhakrishnan, G. V., Van Nieuwerburgh, F., Deforce, D., Chang, C., Karol, K. G., Hedrich, R., Ulvskov, P., Glöckner, G., Delwiche, C. F., Petrásek, J., Van De Peer, Y., Friml, J., Beilby, M. J., Dolan, L., Kohara, Y., Sugano, S., Fujiyama, A., Delaux, P. M., Quint, M., Theißen, G., Hagemann, M., Harholt, J., Dunand, C., Zachgo, S., Langdale, J., Maumus, F., Van Der Straeten, D., Gould, S. B. & Rensing, S. A. 2018. The Chara Genome: Secondary Complexity and Implications for Plant Terrestrialization. Cell 174, 448–464.

Noedoost, F., Sheidai, M., Riahi, H. & Ahmadi, A. 2015. Genetic and morphological diversity in Chara vulgaris L. (Characeae). Acta Biologica Szegediensis, 59, 127–137.

Nordstedt, O. 1889. De Algis et Characeis. Lunds Universitets års-skrift. Lund: Acta Universitatis Lundensis.

Payo, D. A., Leliaert, F., Verbruggen, H., D’hondt, S., Calumpong, H. P. & De Clerck, O. 2013. Extensive cryptic species diversity and fine-scale endemism in the marine red alga Portieria in the Philippines. Proc Biol Sci, 280, 20122660.

Pérez, W., Casanova, M. T., Hall, J. D., Mccourt, R. M. & Karol, K. G. 2017. Phylogenetic congruence of ribosomal operon and plastid gene sequences for the Characeae with an emphasis on Tolypella (Characeae, Charophyceae). Phycologia, 56, 230–237.

Pérez, W., Hall, J. D., Mccourt, R. M. & Karol, K. G. 2014. Phylogeny of North American Tolypella (Charophyceae, Charophyta) based on plastid DNA sequences with a description of Tolypella ramosissima sp. nov. J Phycol, 50, 776–89.

Pérez, W., Hall, J. D., Mccourt, R. M. & Karol, K. G. 2015. OOSPORE DIMENSIONS AND MORPHOLOGY IN NORTH AMERICAN TOLYPELLA (CHAROPHYCEAE, CHAROPHYTA). J. Phycol., 51, 310–320.

Poong, S.-W., Lim, P.-E., Phang, S.-M., Gerung, G. S. & Kawai, H. 2013. Mesospora elongata sp. nov. (Ralfsiales, Phaeophyceae), a new crustose brown algal species from the Indo-Pacific region. Phycologia, 52, 74–81.

Posada, D. & Crandall, K. A. 2002. The effect of recombination on the accuracy of phylogeny estimation. J Mol Evol, 54, 396–402.

Proctor, V. W. 1980. HISTORICAL BIOGEOGRAPHY OF CHARA (CHAROPHYTA): AN APPRAISAL OF THE BRAUN‐WOOD CLASSIFICATION PLUS A FALSIFIABLE ALTERNATIVE FOR FUTURE CONSIDERATION <sup>1</sup>. J. Phycol., 16, 218–233.

Ral Colour System. https://www.ralcolorchart.com/ral-classic/brown-hues x[Online]. [Accessed].

Ray, S., Pekkari, S. & Snoeijs, P. 2001. Oospore dimensions and wall ornamentation patterns in Swedish charophytes. Nordic Journal of Botany, 21.

Ronquist, F. & Huelsenbeck, J. P. 2003. MrBayes 3: Bayesian phylogenetic inference under mixed models. Bioinformatics, 19, 1572–4.

Schaible, R., Gerloff-Elias, A., Colchero, F. & Schubert, H. 2012. Two parthenogenetic populations of Chara canescens differ in their capacity to acclimate to irradiance and salinity. Oecologia, 168, 343–53.

Schneider, S. C., Nowak, P., Von Ammon, U. & Ballot, A. 2016. Species differentiation in the genus Chara (Charophyceae): considerable phenotypic plasticity occurs within homogenous genetic groups. European Journal of Phycology, 51, 282–293.

Schneider, S. C., Rodrigues, A., Moe, T. F. & Ballot, A. 2015. DNA barcoding the genus Chara: molecular evidence recovers fewer taxa than the classical morphological approach. J Phycol, 51, 367–80.

Sheth, B. P. & Thaker, V. S. 2017. DNA barcoding and traditional taxonomy: An integrated approach for biodiversity conservation. Gujarat: Saurashtra University.

Sleith, R. S. & Karol, K. G. 2021. Global high-throughput genotyping of organellar genomes reveals insights into the origin and spread of invasive starry stonewort (Nitellopsis obtusa). Biological Invasions, 23, 3471–3482.

Soulié-Märsche, I. & García, A. 2015. Gyrogonites and oospores, complementary viewpoints to improve the study of the charophytes (Charales). Aquatic Botany, 120, 7–17.

Urbaniak, J. 2003. Tolypella nidifica (O. F. Müll.) A. Braun 1856. Ruggell: A.R.G. Gantner Verlag Kommanditgesellschaft.

Urbaniak, J., Langangen, A. & Van Raam, J. 2012. Oospore Wall Ornamentation in the Genus Tolypella (Charales, Charophyceae). J Phycol, 48, 1538–45.

Van De Weyer, K. & Schmidt, C. 2018. Bestimmungsschlüssel für die aquatischen Makrophyten (Gefäßpflanzen, Armleuchteralgen und Moose) in Deutschland. Band 1, 2. aktualisierte Auflage: Bestimmungsschlüssel [Determination Key for Aquatic Macrophytes (Vascular Plants, Characeaen and Mosses) in Germany. Vol. 1: Determination Key]., Potsdam, Landesamt für Umwelt (LfU) Brandenburg.

Winter, U. & Kirst, G. O. 1991. Vacular sap composition during sexual reproduction and salinity stress in Charophytes. Bull. Soc. bot. Fr., 138, 85–93.

Wood, R. D. 1965. Monograph of the Characeae. In: Wood R.D., I. K. (ed.) A Revision of the Characeae. Weinheim: J Cramer.

